# Improving the detection sensitivity of calcium transients in densely labeled neuronal tissue with pinhole illumination – A low-cost approach

**DOI:** 10.64898/2026.06.18.733097

**Authors:** Caroline Li, Jian-young Wu

**Author notes:** Corresponding author (JW).

## Abstract

Optical recording from large numbers of neurons is an indispensable technique for studying neuronal ensembles. We use optical sectioning through pinhole illumination to reduce the background fluorescence (F_0_) and increase the optical signal (ΔF/F_0_) in ex vivo brain slices densely labeled with GCaMP6f, allowing an ordinary fluorescence microscope to capture calcium transients from over 300 individual CA1 neurons – a marked increase compared to ordinary wide field fluorescence illumination.

Multiple layers of overlapping neurons can be identified by their locations and the shape in space of their ΔF/F_0_ images. A single pinhole mask was placed at the field stop of a wide field illuminator, and the image of the pinhole was projected onto the tissue by a 20X NA 0.95 water immersion objective (Olympus). This created an illuminated disk with a diameter of ∼200 μm and optical sections of hippocampal CA1 pyramidal layer tissue ∼100 μm thick. This illumination blocked a large fraction of the F_0_, which in turn increased the ΔF/F_0_ 5-10-fold compared to that of wide field illumination. When putative pyramidal neurons fire sparsely in the brain slice, up to 300 partially superimposed neurons can be identified by their shape and spatial location in the thick (∼480 μm) ex vivo slice in the CA1 area surrounding the pinhole image. The signal-to-noise ratio was adequate even at a low excitation light level of ∼20k photoelectrons per pixel well on the camera, allowing for 3,000 seconds of total recording time without significant bleaching. This pinhole “half confocal” method has created a useful way to sample calcium transient signals in thick tissue with a large population of neurons densely labeled with GCaMP-6f.

## Introduction

Optical recording of calcium transients from large numbers of neurons is essential for understanding neuronal ensembles during brain processes [1–4]. Optical signals from neuronal activity are fractional changes (ΔF/F_Res_) in fluorescence intensity, where the ΔF is the change in fluorescence caused by neuronal activity, and F_Res_ is the resting fluorescence of the labeled neuron. In densely labeled brain tissue, the F_Res_ from labeled neighboring neurons also contributes to the background fluorescence (F_0_), meaning that F_0_ can be >100 times larger than the F_Res_ of individual neurons. Under wide field fluorescence imaging, the calcium transient signal is represented as ΔF/F_0_ instead of ΔF/ΔF_Res_. ΔF/F_0_ can be as small as 1/100^th^ of ΔF/ΔF_Res_. Such a small ΔF/F_0_ value is often impossible to detect without an effective way of removing F_0_.

Two-photon [5–13], confocal [14–17], and light sheet [18–22, 24] microscopes have been successfully used to record from large numbers of neurons; however, these techniques are often associated with a high instrument cost, making them inaccessible to vast numbers of researchers.

Here, we describe a low-cost, minimally engineered strategy to record large numbers of neurons using an ordinary fluorescence microscope. Following the basic concept of a confocal microscope, pinhole illumination reduces illumination light above and below the focus plane to provide optical sectioning around the focal point [23]. To reduce cost, we employ a fixed pinhole illuminator (“half-confocal”) without a scanner, rather than a full confocal scanning setup. This approach enables ordinary fluorescence microscopes to achieve significantly improved detection performance using a simple filter-like add-on pinhole plate. Furthermore, the lack of a scanning device means that the frame rate can be high (100 frames per second), as it is only limited by that of the camera.

This “half confocal” idea was tested in thick ex vivo brain tissue densely labeled with calcium indicator (thy-promoted GCaMP6f [35]), where, under bright field fluorescence imaging, the ΔF/F_0_ of individual neurons’ calcium transients were about 0.1–1%. The use of an illuminating pinhole focused on the tissue would reduce the F_0_ on the focal plane by 5-10-fold and increase the ΔF/F_0_ by the same order of magnitude. Such an increase in ΔF/F_0_ would allow many more neurons to be identified.

Using pinholes much larger than neurons (200 μm), we identified calcium transients from 300 neurons. In our tests, calcium signals with ΔF/F_0_ values ranging from 1-10% were seen from many neurons surrounding the edge of the pinhole. While the improved ∼1% ΔF/F_0_ is modest, the resulting signal-to-noise ratio is adequate to allow for the identification of neurons using low-cost 12-bit CMOS cameras.

## Methods

### Hippocampal ex vivo slice preparation

P21-P33 male and female C57BL/6J-Tg (Thy1-GCaMP6f) GP5.5Dkim/J. mice (Jax 024276) [35] were used to prepare hippocampal slices in accordance with a protocol approved by the Institutional Animal Care and Use Committee at Georgetown University Medical Center. Following deep isoflurane anesthesia, the animals were rapidly decapitated. The whole brain was subsequently removed and chilled in iced (0-4 °C) sucrose-based artificial cerebrospinal fluid (sACSF) containing (in mM) 252 sucrose; 3 KCl; 2 CaCl_2_; 2 MgSO_4_; 1.25 NaH_2_PO_4_; 26 NaHCO_3_; 10 dextrose; 10 mM HEPES bubbled with 95% O_2_, 5% CO_2_. Hippocampal slices (480 µm thick) were cut in horizontal sections from the dorsal to ventral brain with a vibratome (Leica, VT1000S). Slices were incubated in ACSF for at least 2 hours before each experiment. ACSF used for maintenance and recording contained (in mM) 132 NaCl; 3 KCl; 2 CaCl_2_; 2 MgSO_4_; 1.25 NaH_2_PO_4_; 26 NaHCO_3_; 10 dextrose; 10 mM HEPES bubbled with 95% O_2_, 5% CO_2_ at 26 °C. More detailed methods for brain slices are described in our previous publications [32,33,34,36,38]. Slices were continuously perfused on both sides during imaging. To increase the spontaneous firing rate of the CA1 neurons, in some experiments, 2-20 μM of 4-aminopyridine (4-AP) was added to the perfusion ACSF.

### Imaging

An upright microscope (Olympus BX51 WI) with an epi-illumination arrangement was used. Excitation light was produced by a 470 nm LED (ThorLabs) and an eGFP filter cube (Chroma) with a 425-475 nm excitation filter and a 480 nm dichroic mirror. A 485 nm longpass emission filter was used for the excitation/emission light path. A 20x water immersion objective (0.95 NA, Olympus) was used for the imaging. Pinhole illumination was achieved by closing the iris of the epi-illuminator field stop to the minimum size, projecting an illuminated disk of ∼200 μm onto the tissue. Imaging was conducted using an 80×80 cooled CCD camera (NeuroCCD, Redshirt Imaging-SciMeasure). The camera has a well size of ∼100,000 photoelectrons/well. We adjusted the illumination intensity to ∼50% of the full well inside the illumination disk. The illumination intensity at the brain tissue was <0.1 mW/mm^2^. The imaging frame rate was 100 frames/second. The camera was controlled by the Turbo SM-64 software (Redshirt Imaging-SciMeasure).

Imaging was conducted in intermittent recording trials. The recording trials each lasted 100 seconds, with a 100-second dark resting period between any two imaging trials. The photobleaching was about 2% per 100 seconds of exposure, and the signal-to-noise ratio after ∼3,000 seconds of exposure was still adequate.

### Identifying neurons

Manual analysis was performed using the Turbo SM-64 software (Redshirt Imaging-SciMeasure) in accordance with the following protocol. Through the software, the fluorescence intensity at each pixel was smoothed using 30-frame averages. First, regions of interest (ROIs) were arbitrarily selected around the illuminated pinhole disk (Fig 1B). The fluorescence intensity of the ROIs was plotted through the entire imaging period (Fig 1C, color traces) to identify the peaks of putative calcium transient signals (Fig 1C, 1-31). A background frame either directly before or directly after the peak was then subtracted from the frame at the top of each peak, referred to as the peak time (Fig 1C, white arrows). The subtracted images yield a linear gray-scale image where black and white represent fluorescence intensities between the baseline and the peak, for each pixel (Fig 1D, 1-31). The gray-scale image represents one calcium transient from a putative neuron, which is characterized by the shape and location with respect to the pinhole disk. Next, a neuronal ROI (nROI) is established with all the bright pixels of that calcium transient. Calcium transients with identical nROIs are interpreted as repetitive firing of the same neuron.

**Fig 1.**
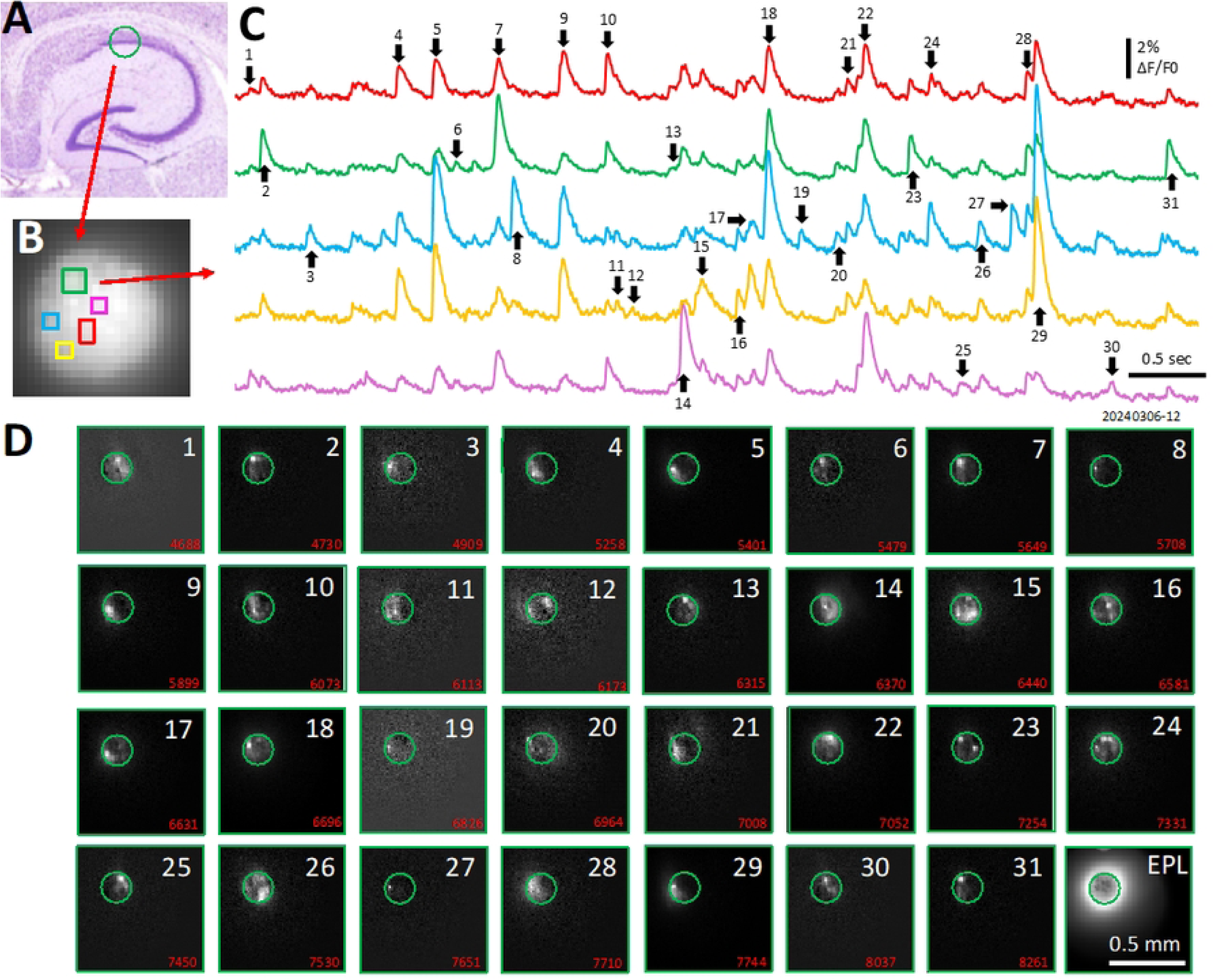
Manual identification of calcium transients using Turbo SM-64 software. **(a)** The CA1 stratum pyramidale layer was imaged by the pinhole. **(b)** Five arbitrary ROIs were selected in the pinhole. **(c)** The ΔF/F_0_ traces of the ROIs were plotted, showing calcium spikes. **(d)** Images were generated by subtracting the image at the ΔF peak, marked by the white arrows in **(b)**, from the baseline image before each peak. **EPL**, an epileptiform event.

### Calculating ΔF/F_0_ of calcium transients

The ΔF/F_0_ of calcium transients was measured within the Turbo SM-64 software (Redshirt Imaging-SciMeasure). The ΔF/F_0_ (%) was defined as ((F_1_-F_0_)/F_0_) * 100% [24], where F_1_ is the averaged fluorescence intensity from the ten brightest pixels of the nROI at the peak time, and F_0_ is the averaged intensity of the same pixel group in the background frames. Calcium transients in our experiments are closely related to the spiking activity of the neurons [25].

## Results

### Large numbers of calcium transients

Calcium transients were mostly seen in the stratum pyramidale of the CA1 subarea (Fig 1A). The ΔF/F_0_ of the transients varied from 1-5% and were clearly distinguished from the baseline noise (Fig 1C, traces). To examine the detection ability of the pinhole illumination method, we created sparse firing conditions in the pyramidal cell population using a low concentration (1-10 μM) of 4-AP in the perfusing bath. Meanwhile, sharp waves initiated from the CA3 were eliminated by low doses of carbachol (0.1-1 μM). With such sparse firing conditions, the calcium transients were distributed evenly in the tissue, at a frequency of 0.2-0.4 transients/second, occurring at different locations in the imaging field.

In a representative recording trial of 100 seconds, we identified 30 calcium transient events at different locations surrounding the pinhole (Fig 1C, 1-31). In this representative preparation, we performed 32 intermittent recording trials, with a total of 3,200 seconds of imaging, leading to the identification of 977 calcium transients. By subtracting the image at the peak of the transient from a background frame, we obtained a gray-scale image of ΔF/F_0_, with brighter pixels representing higher ΔF/F_0_ values. We refer to these images as “spike images” (Fig 1D, 1-31). The locations, shapes, and sizes (number of pixels) of the spike images were used to identify putative neurons. We found that the 977 calcium transients belonged to 322 distinct spiking images (Fig 2), indicating that we identified 322 individual neurons, or branches of neurons.

**Fig 2.**
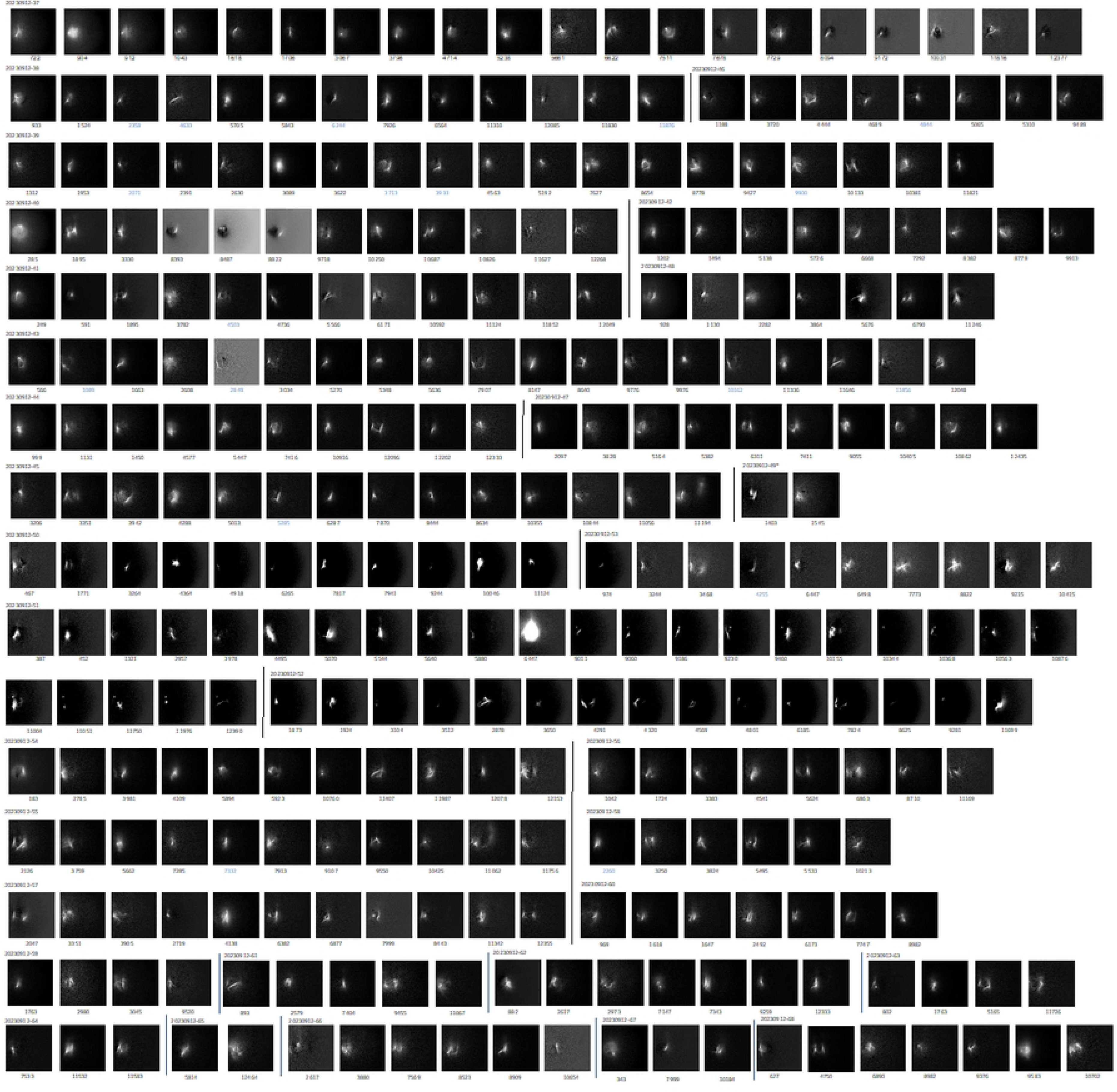
Identification of individual neurons from calcium transient imaging. A representative recording showing calcium transients detected over 3200 seconds of imaging in the CA1 stratum pyramidale. Individual neurons were identified based on the size, shape, and spatial location of their fluorescence signals, yielding 322 putative neurons from 977 calcium transients.

Calcium signals from different neurons often overlap in a singular location. For example, 26 of the 31 transients can be observed in the same ROI (Fig 1C, red trace). This indicates that we can detect signals from superimposed soma or branches of neurons. This capability is distinct from two-photon or confocal microscopy, where only a thin section of neurons can be visualized at a time.

During the entire recording period (3,200 seconds), the majority of the spike images (196 out of 322) were observed only once. While a few spike images occurred repeatedly, only 19 of them were observed 10 times or more (Fig 3). This pattern of sparse firing largely reduced the likelihood of coincidence, *i.e.,* two neurons fired concurrently and were mistakenly identified as a single new spike image.

**Fig 3.**
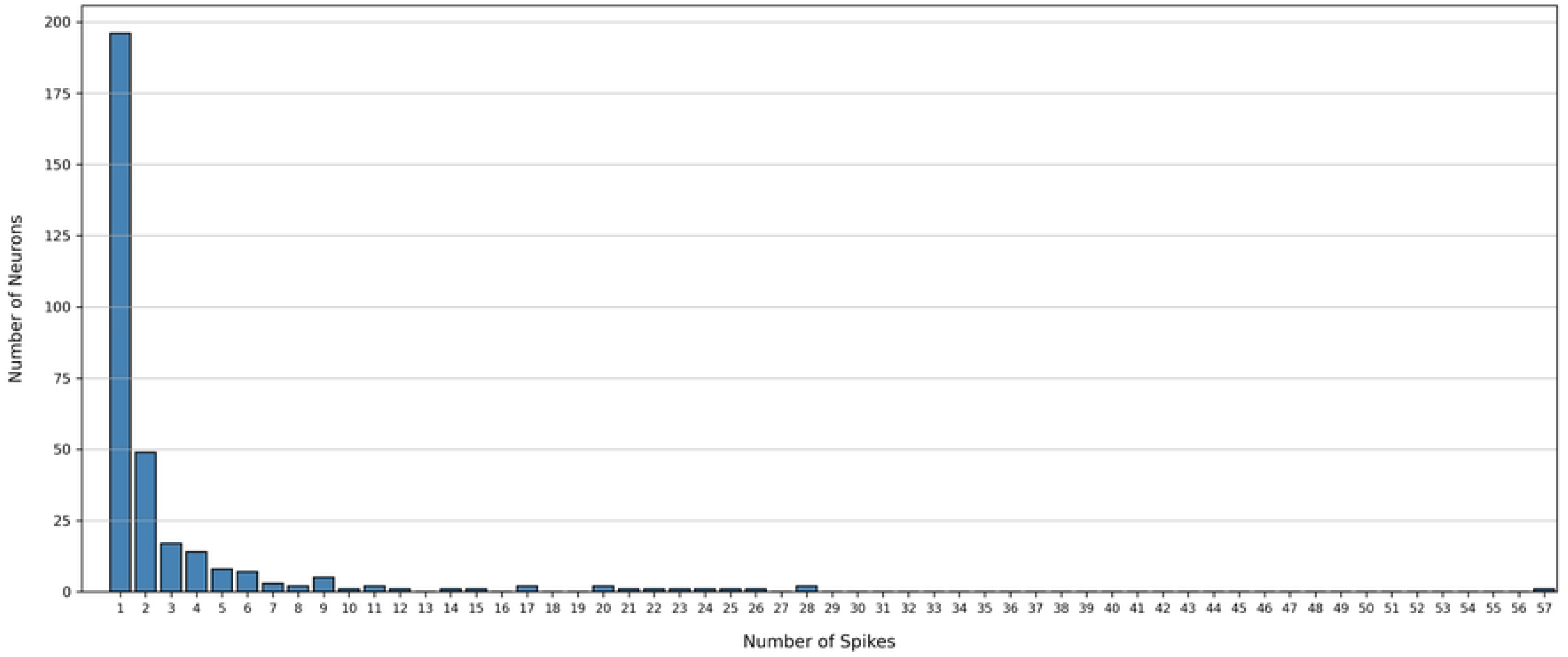
Spike count of the 322 neurons identified in. Fig 2 Most of the neurons (196 out of 322) fired only once, while a few fired more frequently. One neuron fired 57 times over the 3200-second-long imaging period.

Spike images with a similar shape, size, and location were frequently seen (Fig 2), and these similar images were carefully checked by manual inspection of the outlines of the bright pixels corresponding to each spike image (Fig 4A). Then, the outlines were superimposed, allowing for the differentiation of the size and location of the neurons (Fig 4B). Spike images with minor differences in size and location were removed from the neuronal catalog in Fig 2. Superimposing the outlines was particularly useful for examining neurons where the spike images had no distinguishing morphological characteristics (e.g., when there were no branching features).

**Fig 4.**
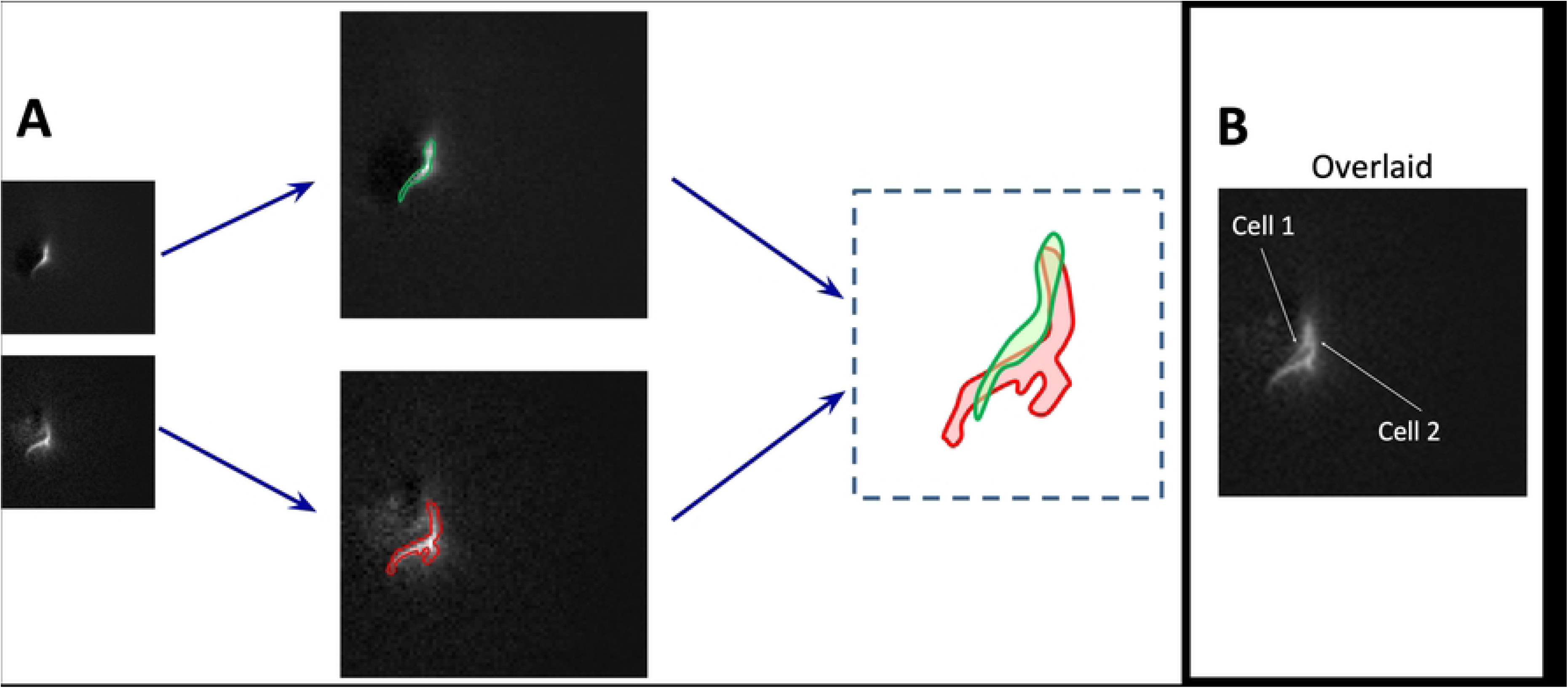
Identification of highly overlapped neurons. **(a)** Two neurons that have a similar shape and spatial location appear as one neuron. However, the ambiguity can be solved by overlaying the outlines of the two images of the neurons (red and green lines). **(b)** Overlapping the images of the two neurons helps emphasize subtle differences between them.

In a fully analyzed preparation, 845 calcium transient spikes were identified in one imaging field. Considering the sparseness of firing, we estimate that approximately 100-300 neurons can be identified in the imaging field. Across 4 additional analyzed preparations with a total imaging time of 3,200 seconds and spanning 4 imaging fields, we identified 1,228 spikes. From one of these imaging fields, we identified around 80-100 neurons during a 1,000-second imaging period. The ΔF/F_0_ values of these spikes were around 1-5%, well above the background imaging noise. This suggested that under conditions of sparse firing and extended imaging durations, around 300 neurons can be identified in an imaging field.

### Comparison with wide field illumination

From the same tissue and field of view, we compared the calcium transients found in wide field and pinhole illuminations. Under wide field illumination, fewer calcium transients were distinguishable from the baseline noise, suggesting that reducing the background fluorescence F_0_ can significantly improve the detection ability.

Additionally, the difference in ΔF/F_0_ was significant. In one data set, wide field and pinhole illumination were performed in the same field of view, where 30 neurons (123 firing events) appeared in both wide field and pinhole trials. The mean ΔF/F_0_ was 4.43% in pinhole illumination trials and 1.57% in wide field illumination trials, demonstrating a ∼3-fold increase in ΔF/F_0_. The amplitude distributions of the calcium transients recorded from the two illumination methods showed a marked difference, where a large fraction recorded with wide field illuminations had a low amplitude (Fig 5A). The ΔF/F_0_ obtained from pinhole illumination trials exhibited greater variability (variance = 5.00), as compared to that of wide field illumination trials (variance = 1.02). Lower variability suggests that some ΔF/F_0_ transients were not detected in the wide field illumination trials. Overall, the ΔF/F_0_ from the two illumination methods were significantly different (p < 0.001) in a two-tailed Welch’s t-test (df = 89, Fig 5B).

**Fig 5.**
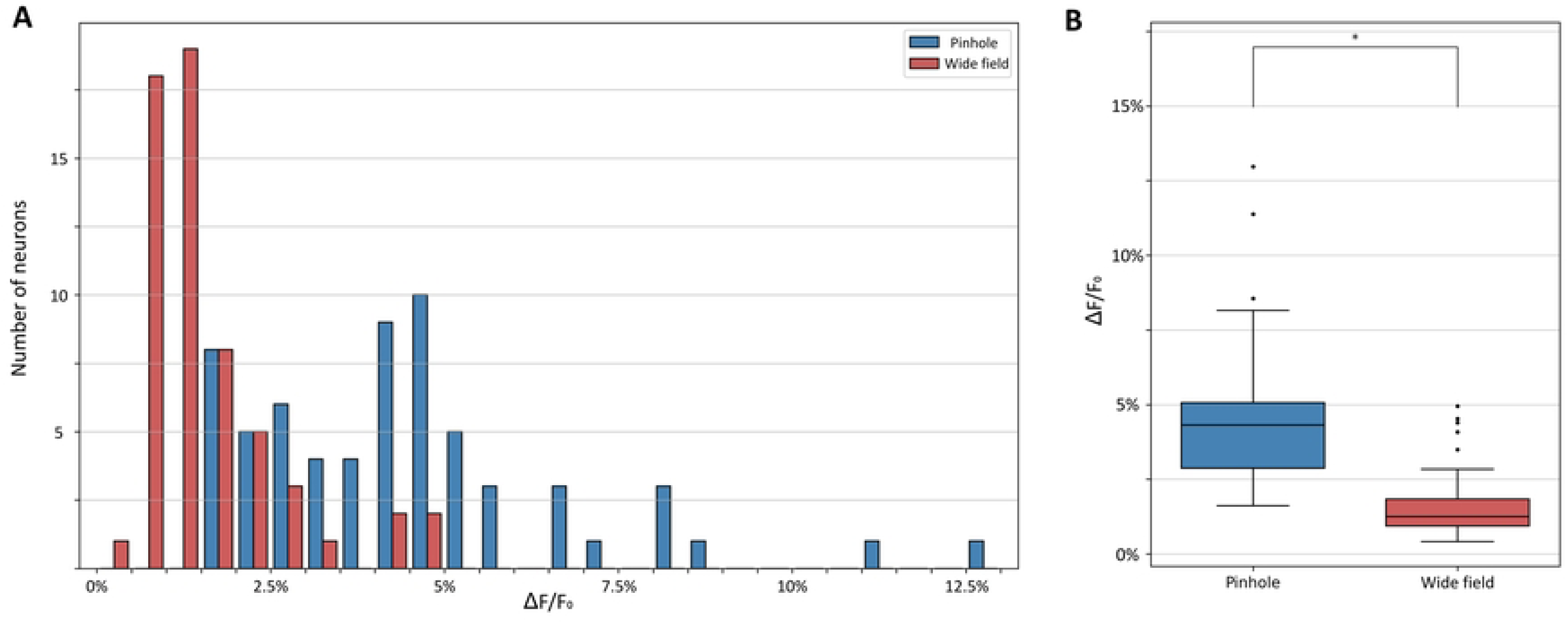
Comparison with wide field illumination. **(a)** Neurons identified from the same tissue with pinhole or wide field illuminations. **(b)** ΔF/F_0_ of pinhole or wide field illumination.

In another preparation, we cataloged the neurons of both wide field and pinhole illumination trials. Differences in the number of calcium transient spikes, unique neurons, and ΔF/F_0_ values were revealed. Out of 319 spike images identified from the pinhole illumination trials, 128 were classified as unique neurons (Fig 6A). In contrast, within the same field of view, only 52 spike images were identified from the wide field illumination trials, with 37 being classified as unique neurons (Fig 6B). The mean ΔF/F_0_ value for the wide field trials was 0.96% and the mean ΔF/F_0_ for the pinhole trials was 2.03% with a significant (p < 0.001) difference in a two-tailed Welch’s t-test (df = 366) (Fig 6C, right panel). The variance in the ΔF/F_0_ distribution was also different (pinhole variance =12.85, wide field variance = 0.422).

**Fig 6.**
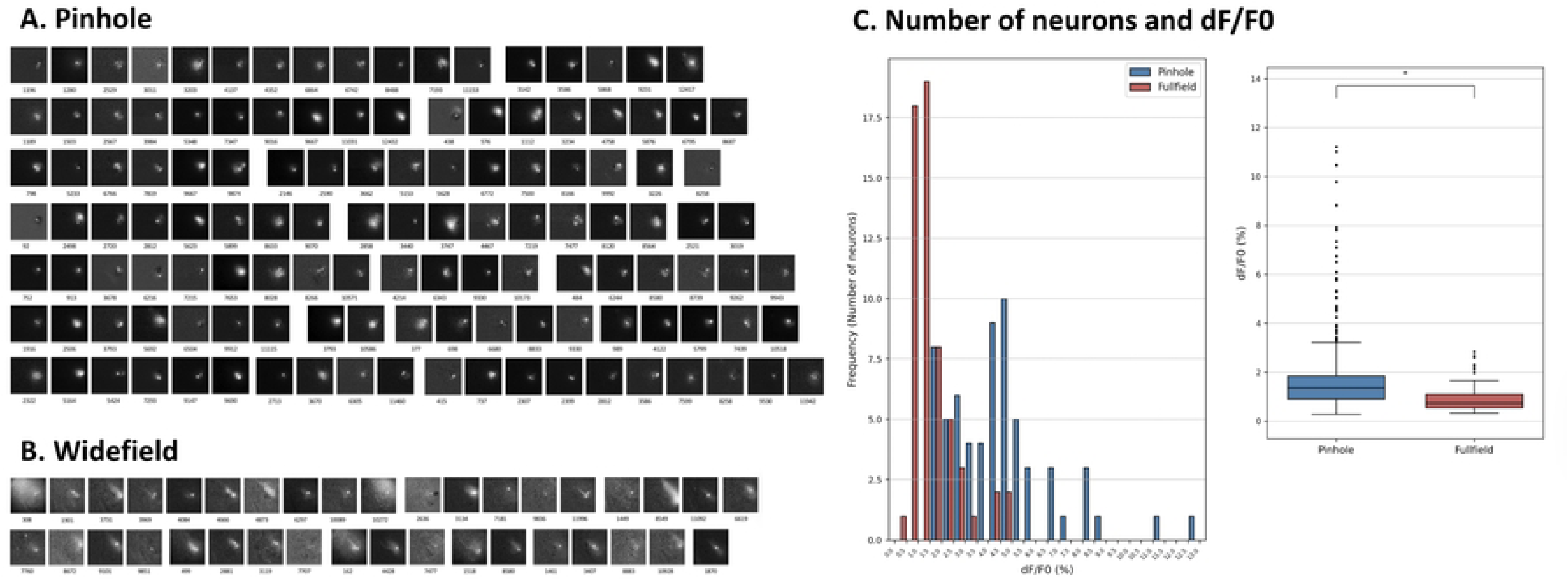
Neurons identified with pinhole or wide field illumination. **(a)** Images of the neurons identified through pinhole illumination. **(b)** A fraction of the same neurons identified through wide field illumination. **(c)** Comparison of the number of neurons and ΔF/F_0_ between pinhole illumination and wide field illumination.

### Bright ring during epileptiform events

To further illustrate the enhancement of the ΔF/F_0_ through pinhole illumination, we measured the calcium transients of epileptiform events. Spontaneous epileptiform events (interictal-like spikes, IS) occasionally occurred in our ex vivo hippocampal slices. In our bath containing a low concentration of 4- AP (∼2 μM), the occurrence rate of IS increased, often appearing once every few hundred seconds during imaging. The calcium transient signals associated with IS exhibit high amplitudes (ΔF/F_0_ of 100-500%) and long durations (> 1 second), markedly different from calcium transients of individual spikes (Fig 7). These high amplitudes and long durations resulted from the spikes and bursts of a large fraction of neurons in the tissue during an IS.

**Fig 7.**
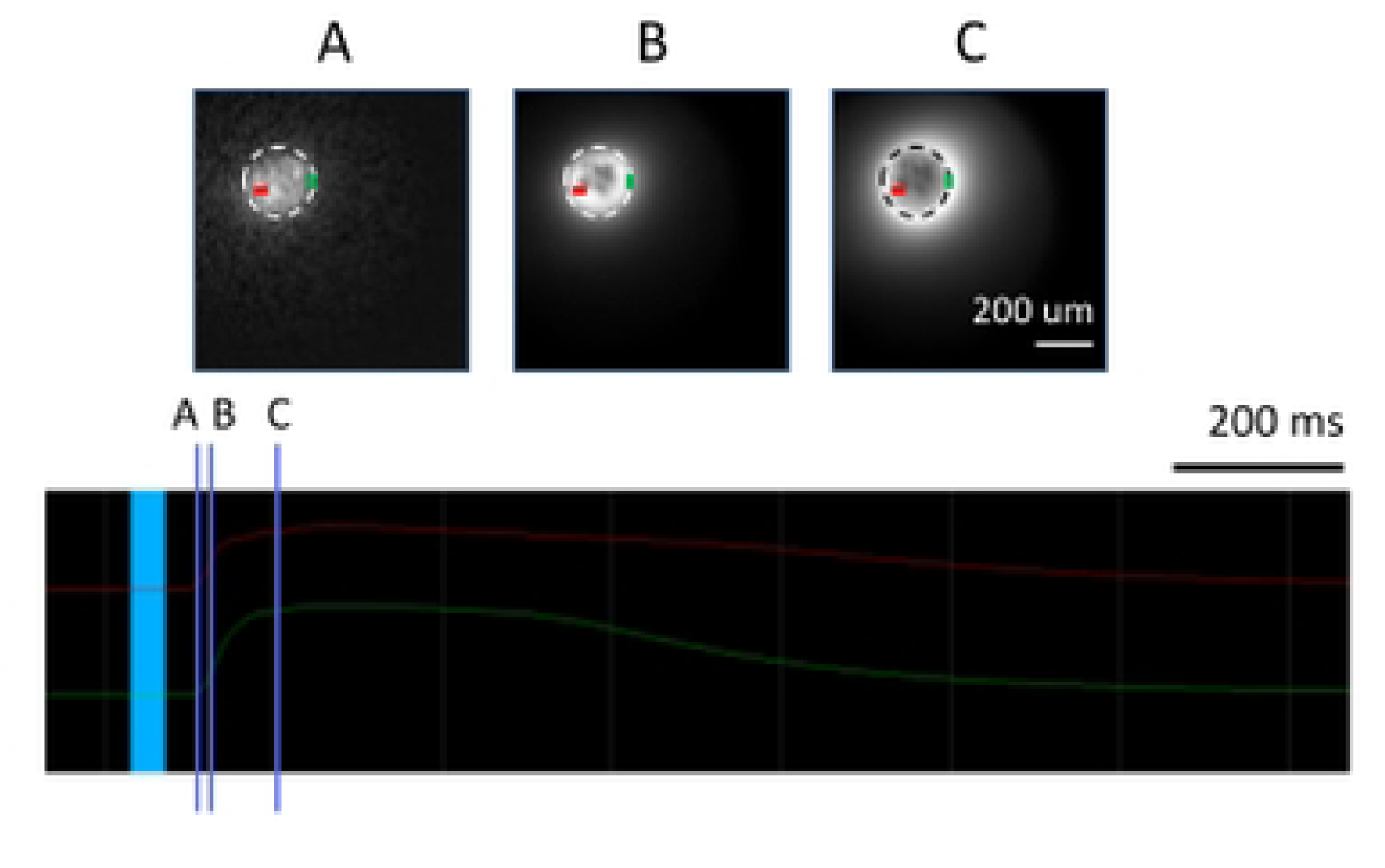
The edge effect of pinhole illumination. The top images (A-C) are subtracted images at the onset of an epileptiform event. The timing of the subtraction is labeled by the three vertical blue lines, A, B, and C, on the bottom traces. The background level for the subtraction is labeled by the light blue area on the bottom traces. Bottom traces are calcium transients from two areas (red and green, labeled in the top images). A larger ΔF/F_0_ was seen surrounding the edge of the pinhole (marked by broken line circles). Note that at different times of the event, higher ΔF/F_0_ values occurred at the inner and outer edges of the pinhole.

After background subtraction, the images of epileptiform events have a bright annulus around the edge of the pinhole (Fig 7B), demonstrating an enhanced ΔF/F_0_ surrounding the edge of the pinhole.

Fig 7 also shows subtracted images from three time points during the rising phase of an IS (top panel A-C) in reference to the amplitude plot (bottom panel). At the early initial phase, a few bright spots occurred in the subtracted image (Fig 7, panel A), suggesting the activity of early-firing neurons before most neurons were activated. Later, a bright annulus occurred surrounding the image of the pinhole (panels B and C), suggesting an enhanced ΔF/F_0_ near the edge of the pinhole. While individual neurons cannot be detected during IS, the edge effect of the pinhole illumination was clearly seen.

### Multiple pinholes

We also used multiple pinholes to expand the imaging field and increase the number of neurons that could be detected. In an experiment shown in Fig 8, we used 5 additional pinholes surrounding the central pinhole. Neurons detected in the peripheral pinholes have similar ΔF/F_0_ values (Fig 8B), suggesting that multiple pinholes are effective and can capture more neurons.

**Fig 8.**
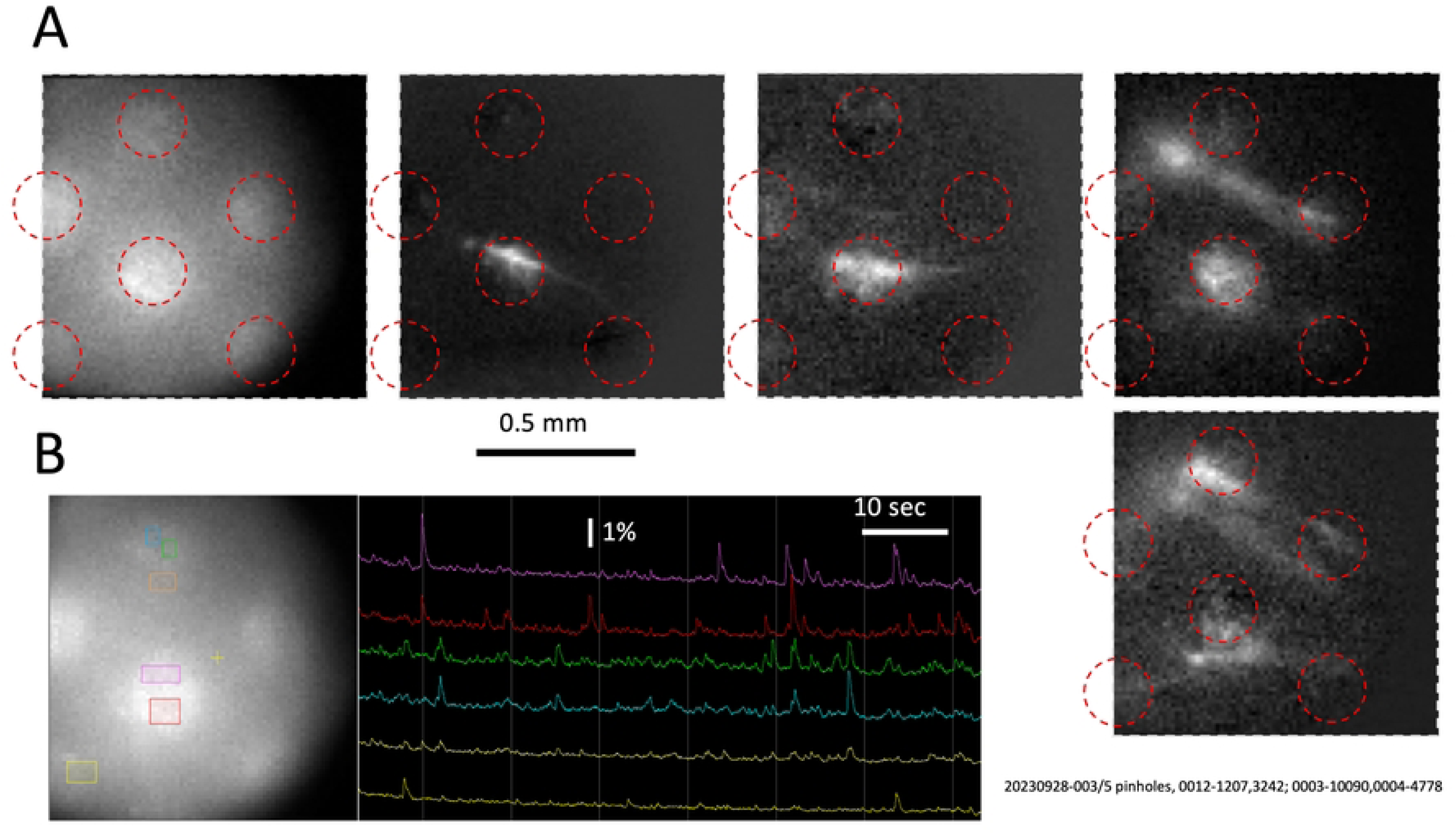
Neurons identified with multiple pinholes. **(a)** Five pinholes (marked by red broken lines) captured neurons from a greater area. Note that neurons could be seen even in the tissue outside of a pinhole. **(b)** Traces showing neuronal activity in multiple pinhole illumination.

## Discussion

The pinhole illumination method described in this report aims to enhance the detection capability of wide-field illumination without significantly increasing the cost. By incorporating a pinhole mask at the field diaphragm of the epi-illuminator, the detection sensitivity increased by a factor of 5-10 (Fig 5). Increased detection capability was reflected by an increased ΔF/F_0_ of the same neuron between pinhole and wide field illuminations (Fig 5D).

From the same tissue, many more neurons can be detected through pinhole illumination (Fig 6A, B) compared to wide field illumination, suggesting that pinhole illumination boosts calcium transient signals into the detectable range; neurons that have low ΔF/F0s were detected through pinhole illumination but are more likely to be missed in wide field illumination (Fig 6C). In one example shown in Fig 2, we captured 977 calcium transients from tissue with sparse firing and identified 322 putative neurons by their distinct shapes. This number is comparable to the hundreds of neurons that Li et al. was able to scan by standard-two photon microscopes in a 300 x 300 µm field of view [39]. Additionally, a two-photon in vivo imaging study conducted within a 500 µm-wide cortical column by Kaszas et al. reported the detection of 51–291 neurons [40]. Within our 200 μm pinhole, we were able to detect a greater number of neurons through pinhole illumination.

The ΔF/F_0_ values observed in our experiments generally ranged from 1–5%, substantially lower than the 100–500% signals commonly reported for two-photon microscopy studies [5,10]. However, the recordings also exhibited lower noise levels due to the high optical throughput and reduced shot noise associated with single-photon excitation. Consequently, the detected calcium transients maintained ΔF/F_0_ amplitudes well above baseline noise levels (Fig 1C). Notably, a ΔF/F_0_ threshold of approximately 6% has previously been shown to be sufficient for classifying visually responsive neurons in two-photon imaging experiments conducted by Chen et al [41]. The majority of ΔF/F_0_ values obtained using pinhole illumination came close to meeting this benchmark. Li et al. examined the ΔF/F0 of sharp-wave ripple events using both wide field illumination and two-photon imaging. They found that the average ΔF/F0 of events recorded using wide field illumination was 0.36%, and that the average of events recorded using two-photon microscopes ranged from 30-2000% [32]. Our average ΔF/F0 of 4.43% using pinhole illumination demonstrates a marked improvement from wide field illumination, which contributed to our ability to detect a greater number of neurons.

The pinholes we used were much larger than the putative neurons detected (Figs 1 and 2). Large pinholes can still effectively reduce F_0_, because the reduction of F_0_ is provided by the edge of the pinhole. The edge effect is further verified by the images of epileptic events, where a large fraction of neurons have simultaneous calcium transients. While individual neurons cannot be seen, a bright annulus was seen at the edge of the pinhole, reflecting a higher ΔF/F_0_ associated with the edge (Fig 7). Using large pinhole illumination allows for a field of depth of up to 200 µm, allowing multiple layers of overlapped neurons to be recorded. As shown in Figs 2 and 4, many neurons overlap in space, suggesting that they are from different planes.

With the improved detection capability, we estimate that an ordinary 10-bit CMOS camera (e.g., Thorlabs CS135MU, priced around $2,000) should be sufficient to detect fluorescence changes as small as 0.1% ΔF/F_0_ when signals from more than eight pixels are binned together as a single region of interest (ROI). The read noise of these CMOS cameras is usually negligible (<20 e⁻) compared to the shot noise (∼300 e⁻) generated by typical wide field fluorescence intensity (∼1,000 photoelectrons per pixel well). Using an ordinary 10-bit CMOS camera instead of an EMCCD or high-grade scientific CMOS camera can greatly reduce the cost. Single-photon excitation generates high emission light throughput, allowing for the use of ordinary LEDs as the excitation source, avoiding the need for expensive lasers.

Our design does not require scanning, thereby removing the bottleneck imposed by scanner speed on frame rate. In this report, we used a frame rate of 100 frames per second. The 10-bit camera mentioned above can reach up to 520 frames per second at 640 × 512 resolution, enabling the detection of spiking sequences when fast calcium indicators (e.g., jGCaMP8f [26,27]) are used. Eliminating mechanical scanning and its associated optical components further reduces system complexity and instrument cost.

### How pinhole illumination improves F/F_0_

When a pinhole is added to the field diaphragm of the epi-illuminator, its image is projected to the tissue (Fig 9A), and the neurons outside of the pinhole image are not illuminated, meaning that they do not contribute to F_0_. Neurons at the edge of the pinhole (Fig 9A, red spot) are fully illuminated, but the scattered F_0_ from other neurons is largely reduced. On the other hand, in wide field illumination (Fig 9B), all neurons in the field are evenly illuminated and their resting fluorescence F_0_ is scattered throughout the tissue, reducing the ΔF/F_0_ of active neurons.

**Fig 9.**
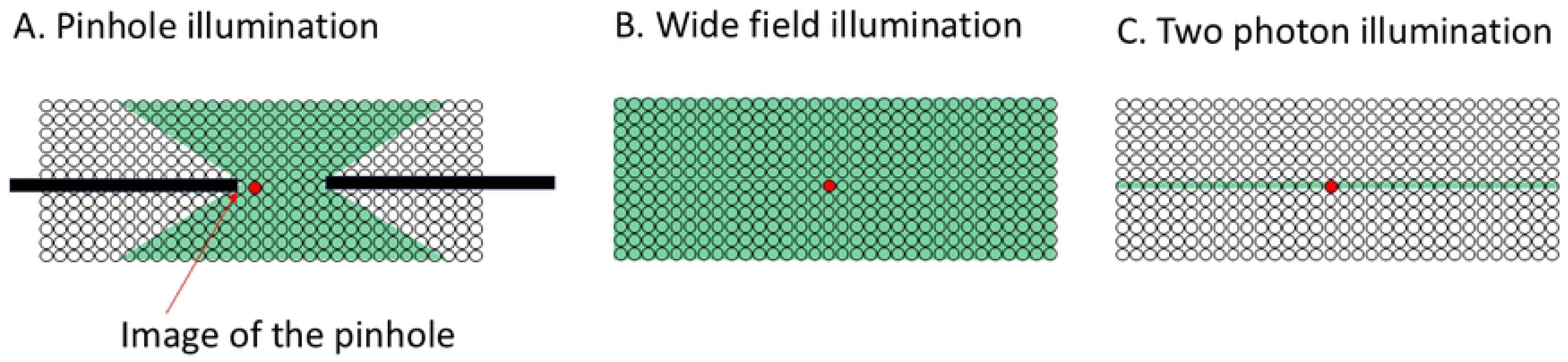
The image of a pinhole reduces F_0_. **(a)** The image of a pinhole (black plate) is projected onto the tissue by a cone of light (green). Neurons outside the light cone (blank circles) do not fluoresce. Thus, the overall F_0_ of the active neuron (red circle) is lower with pinhole illumination compared to wide field illumination, depicted in **(b)**, where all neurons (green circles) produce F_0_. **(c)** Two-photon imaging illuminates only neurons in its focal plane, producing a low F_0_ and high ΔF/F_0_.

The reduction of F_0_ by our pinhole illumination method can be roughly estimated (without considering the difference in refractive index between water and cortical tissue). The image of the pinhole is projected to the tissue by a cone of light (Fig 9A, green region). Only the neurons in the cone are illuminated and contribute to the F_0_. Using the 20X 0.95 NA objective, the pinhole image is projected onto the tissue with an oblique angle of ∼43 degrees. With a working distance of ∼1 mm, the out-of-focus light at 100 μm above or below the 200 μm pinhole should be ∼1/4. Such a 4-fold reduction in illumination should translate to a 1/4^th^ reduction in F_0_ for out-of-focus neurons, and a 4× gain in ΔF/F_0_ for the neurons in the same plane as the pinhole image.

From this estimation, smaller pinholes should experience a larger reduction in F_0_. As shown in Fig 9C, a 100 μm pinhole would get a ∼9-fold reduction in F_0_ at 100 μm off the focal plane, and a 50 μm pinhole should get ∼25-fold reduction. However, the opaqueness of the cortical tissue would greatly reduce the gain of smaller pinholes. Though we have tested smaller pinholes (data not shown), the number of neurons detected did not exceed that of the 200 μm pinhole, probably because smaller pinholes have a shorter edge line and a thinner optical sectioning plate. When using thick tissue slices of ∼500 μm in our experiments, 200 μm pinholes appear to be the most satisfactory.

While the 1-5% ΔF/F_0_ values from our experiments are much smaller than the 100-500% ΔF/F_0_ values obtained from two-photon imaging (Fig 9C), our method, single-photon illumination, offers ignorable shot noise and a much lower instrument cost.

### Limitations and future directions

Our method also has potential limitations. First, the pinhole enhances fluorescence contrast primarily at its edge, meaning that only neurons near the pinhole boundary can be effectively detected. When using a 20× objective, the full imaging field is approximately 1 mm in diameter, while a 200 µm pinhole covers only about 1/25 of that area.

This limitation can be partially addressed by using multiple pinholes (Fig 8). Arranging six pinholes around a central one in a hexagonal pattern would allow for the visualization of ∼7× more neurons within the field of view. However, each pinhole must be separated by a dark surrounding band, as the reduction in F_0_ occurs only at the dark–light interface.

In practice, spike signals are often visible even in the dark regions between pinholes (Fig 8), suggesting that these areas are illuminated by scattered light and that the resulting ΔF/F_0_ remains sufficient for neuron identification.

As a future direction, a fiber-optic–based illuminator could overcome the limitations of pinhole illumination and achieve true full-field illumination. In this design, each pinhole is replaced by the end aperture of an optical fiber, and the light emerging from the fiber serves as the pinhole illumination source. The illuminator consists of a coherent fiber bundle containing hundreds of individual optical fibers (Fig 10). These fibers are organized into four groups, enabling four interlaced illumination patterns.

**Fig 10.**
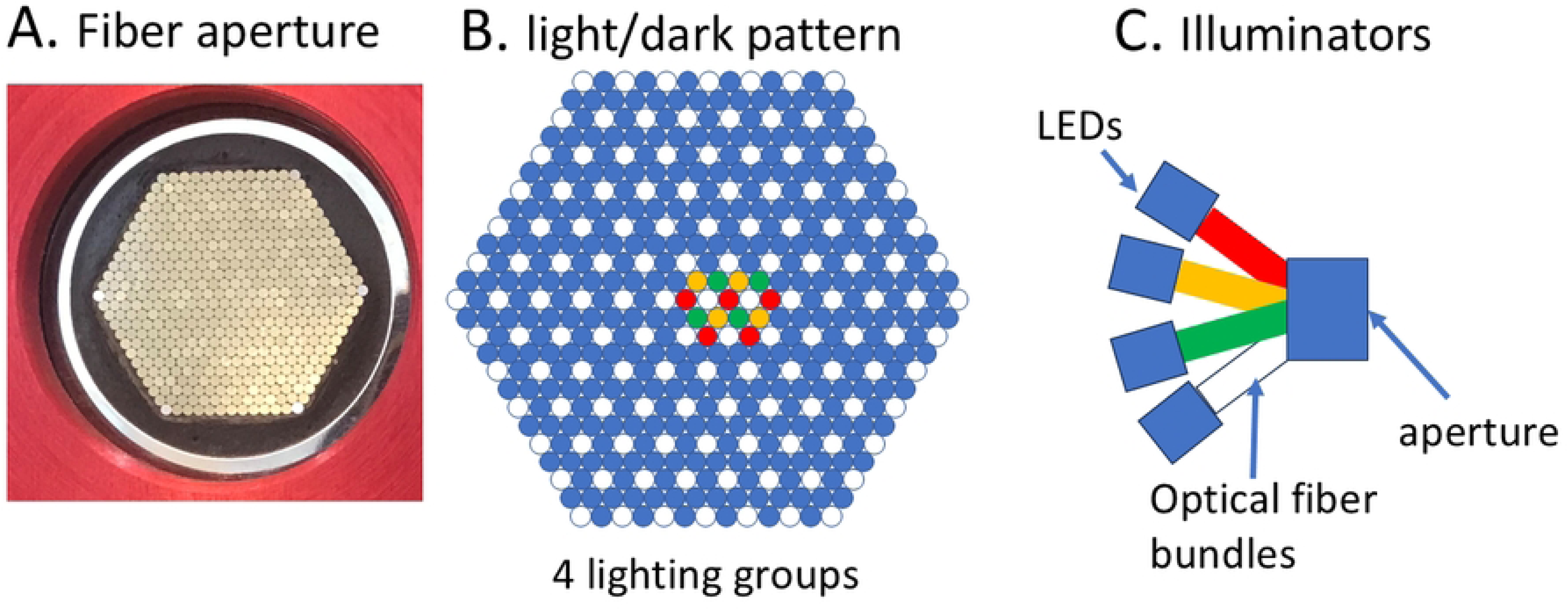
Full field illumination with multiple pinholes. **(a)** A fiber optical bundle is used for illumination, where each fiber serves as a pinhole. **(b)** Optical fibers are grouped into four lighting groups (marked by yellow, green, white, and red). The four lighting groups take turning on, creating an illumination with multiple pinholes occupying the full field, while each pinhole is surrounded by a dark annulus. **(c)** The illuminator. Four LED lights sequentially illuminate the fiber groups (red, yellow, green, and white) to create a virtual Petráň disk without moving parts.

Multiple illumination pinholes would be activated simultaneously and electronically scanned across the tissue, similar in concept to a Petráň multiple-pinhole spinning disk [28–31], but without any moving parts. While this approach would represent an ideal solution for improving ΔF/F_0_ using pinhole illumination, the instrument cost would increase, as a custom illuminator would need to be designed and integrated with the existing fluorescence microscope.

Another limitation of the method is the time-consuming nature of manual data analysis. Our optical section is approximately 100 µm thick, causing neurons that overlap along the z-axis to project onto the same location. Although the thick optical section greatly increases the total number of identifiable neurons, automated shape-recognition algorithms [37] perform poorly in distinguishing overlapping cells. We devoted more than 200 hours of manual analysis to our dataset (imaging of four tissue slices), making extensive manual processing a major limitation to the scalability of this method. In addition, manual analysis introduces individual bias. Artificial intelligence (AI)-based image recognition will eventually alleviate the burden of such intensive manual analysis.

In conclusion, a pinhole mask with 1-7 pinholes provides an inexpensive way to increase detection ability in ordinary fluorescence microscopes. Fiber optics illuminators may further improve this approach, allowing for the identification of even more neurons from an imaging field.

